# Kinetic control in amyloid polymorphism: Different agitation and solution conditions promote distinct amyloid polymorphs of alpha-synuclein

**DOI:** 10.1101/2022.11.25.517910

**Authors:** Santosh Devi, Dushyant Kumar Garg, Rajiv Bhat

## Abstract

Aggregation of neuronal protein α-synuclein is implicated in synucleinopathies, including Parkinson’s disease. Despite abundant in vitro studies, the mechanism of α-synuclein assembly process remains ambiguous. In this work, α-synuclein aggregation was induced by its constant mixing in two separate modes, either by agitation in a 96-well microplate reader (MP) or in microcentrifuge tubes using a shaker incubator (SI). Aggregation in both modes occurred through a sigmoidal growth pattern with a well-defined lag, growth, and saturation phase. The end-stage MP- and SI-derived aggregates displayed distinct differences in morphological, biochemical, and spectral signatures as discerned through AFM, proteinase-K digestion, FTIR, Raman, and CD spectroscopy. The MP-derived aggregates showed irregular morphology with a significant random coil conformation, contrary to SI-derived aggregates, which showed typical β-sheet fibrillar structures. The end-stage MP aggregates convert to β-rich SI-like aggregates upon 1) seeding with SI-derived aggregates and 2) agitating in SI. We conclude that end-stage MP aggregates were in a kinetically trapped conformation, whose kinetic barrier was bypassed upon either seeding by SI-derived fibrils or shaking in SI. We further show that MP-derived aggregates that form in the presence of sorbitol, an osmolyte, displayed a β-rich signature, indicating that the preferential exclusion effect of osmolytes helped overcome the kinetic barrier. Our findings help in unravelling the kinetic origin of different α-synuclein aggregated polymorphs (strains) that encode diverse variants of synucleinopathies. We demonstrate that kinetic control shapes the polymorphic landscape of α-synuclein aggregates, both through de novo generation of polymorphs, and by their interconversion.

## 1. Introduction

Human α-synuclein (α-Syn) is a neuronal protein, which under normal conditions, localizes to the presynaptic terminal of neurons, and is involved in synaptic transmission and exocytosis [1, 2]. A recent study has pointed out its role in modulating processing bodies (P-bodies) dynamics, wherein α-Syn interacts with different RNA-binding proteins of P-bodies and plays an essential part in RNA metabolism [3]. Apart from its function in the brain, α-Syn is reported to be a prominent player in the innate immune response against viral and bacterial infection in the gut [4, 5]. Under unknown circumstances, α-Syn forms amyloid aggregates in the cytoplasm of neurons and accumulates in the form of filamentous cross-β-rich inclusions, also called Lewy bodies-a hallmark of synucleinopathies such as Parkinson’s disease and dementia with Lewy bodies [6, 7]. α-Syn is a ∼14.5 kDa intrinsically disordered protein and consists of three central regions (i) an N-terminal positively charged stretch (residue 1-60), (ii) a hydrophobic non-amyloid-β stretch (NAC region; residue 61-95), which forms the core of α-Syn amyloid, and (iii) a C-terminal acidic tail, region 96-140 that binds to metal ions [8, 9]. Due to the prevalence of oppositely charged amino acid residues at the N- and C-terminal of α-Syn, a long-range transient interaction prevails. Due to this, the average monomeric size of α-Syn is relatively smaller than the equivalent typical random coil of similar size [10–12]. It is due to same reason; a high nucleation barrier exists for in vitro aggregation of α-Syn at pH 7.4. This fact is evident from the observations that in quiescent conditions, it does not undergo aggregation even after several days of incubation and thus requires a high agitation. Due to its structural plasticity, several conformational states of soluble α-Syn exist in the cell, which range from disordered monomers to both disordered and ordered oligomers; and cellular factors are believed to shape those states [12, 13].

The α-Syn conformational state in the cell is dictated by myriad factors such as binding to the membranes and metal ions, interaction with chaperons, post-translational modifications, proteolysis, and interaction with neurotransmitters [13–19]. Yet, the clarity on the molecular mechanism through which these factors play a role in its transition from random coil monomers to β-rich aggregates is lacking. Over the years, many *in vitro* studies have been carried out on recombinantly purified α-Syn, which has uncovered important clues into its aggregation. While test tube conditions do not replicate the actual cellular environment in which α-Syn is present, yet it facilitates studying the role of various cellular factors, and is a bottom-up approach to constructing the overall α-Syn aggregation pathway. A stepwise delineation of the events leading to the initiation and progression of α-Syn assembly under different conditions using different structural probes has unravelled various structural intermediates that populate en route to its aggregation [20–22].

The in vitro aggregation mechanism of α-Syn remains ambiguous with different models reported for its aggregation. The nucleation-dependent polymerization model suggests that the soluble random structure monomers of α-Syn structurally convert to a β-sheet rich species (a rate-limiting step), which interact among themselves and form β-sheet rich oligomers [23]. These oligomers eventually assemble in small fibrils, which further recruit soluble monomers to elongate. Alternatively, the nucleation-conversion-polymerization model posits that random coil monomers undergo intra- and inter-molecular collapse to form disordered oligomers, which slowly restructure into β-sheet-rich oligomers before assembling into the fibrils [24–27]. At neutral pH, α-Syn fibrillation occurs through primary nucleation, whereby de novo conversion of monomers into fibrils occurs and further amplification occurs through the addition of monomers to fibril ends [21]. At slightly acidic pH, in addition to primary nucleation, fibril amplification also occurs through the secondary nucleation, whereby, in addition to fibril ends, fibril surface acts as nucleation sites for aggregation [28, 29]. This difference in the microscopic behavior among primary and secondary nucleation is manifested in the macroscopic kinetic curves of α-Syn aggregation. Recent reports highlight liquid-liquid phase separation as one of the nucleation mechanisms of α-Syn aggregation, wherein α-Syn monomers form highly concentrated liquid-like assemblies driven by its NAC domain [30–33]. Under abnormal circumstances, these liquid assemblies mature into solid aggregates through homogenous primary nucleation [30]. Thus, scope exists for studying the aggregation of α-Syn in different physicochemical conditions.

In this work, we studied the *in vitro* aggregation behavior of α-Syn. We monitored the kinetics of aggregation through thioflavin-T fluorescence vis-à-vis accompanying structural and morphological changes through CD, FTIR, and AFM. We carried out aggregation separately in two commonly used agitation modes - in 96 well plates agitated on a microplate reader (MP); in a microcentrifuge tube agitated on a shaker incubator (SI). We correlated the thioflavin-T monitored progression of α-Syn aggregation with secondary structural change through CD and observed that in the two conditions, the far-UV CD monitored conformational conversion was not concomitant with the thioflavin-T monitored aggregation. The conformational conversion in both the conditions continues to occur after saturation of thioflavin-T kinetics. The end-stage MP incubated aggregates were kinetically arrested in predominantly random conformation, whereas SI incubated aggregates were β-sheet-rich. The MP aggregates convert to β-sheet rich aggregates upon further incubation in SI. Moreover, the kinetic barrier of MP aggregation was also overcome upon either seeding with β-sheet rich SI aggregates or in the presence of cellular osmolytes. Thus, we present two different aggregation routes for α-Syn aggregation that gives rise to two kinetically separated amyloid polymorphs. We also discuss our observations with recently discovered α-Syn aggregated polymorphs exhibiting different sensitivities to thioflavin-T. We further discuss our findings in the context of kinetic barriers shaping the aggregation landscape of the α-Syn that would have implications towards designing effective small-molecule inhibitors for further screening.

## 2. Results

### 2.1. Secondary structure acquisition of α-Syn was not concurrent with the formation of ThT-positive species

We first monitored the aggregation kinetics of α-Syn by measuring its time-dependent ThT fluorescence. In the case of the microplate, we observed that both the lag phase and amplitude of the saturation phase (ThT fluorescence intensity at the final plateau) were highly dependent on the agitation speed (Fig. S2). With an increase in agitation speed, the reduction in the lag phase, and enhancement in the ThT amplitude was evident. Therefore, we chose 900 RPM agitation speed for aggregation to bring the overall kinetic duration within 24 h (Fig. S2). The kinetics in both the agitation modes displayed a sigmoidal pattern with well-defined lag, growth, and saturation phases. The lag time of MP incubated α-Syn was 4.5 h (Fig. 2A; Table 1), and that of SI incubated aggregate was 6.1 h (Fig. 2B; Table 1). The transition midpoint (t_1/2_), a time point where 50% of the kinetics completes was ∼ 6 h for MP incubated α-Syn, and 7.9 h for SI incubated α-Syn. The ThT signal was saturated at 10 h in MP and 12.8 h in SI. We further derived the relationship between t_1/2_ of aggregation kinetics and the initial monomer concentration by plotting both the quantities on a double logarithmic scale. The slope of this plot (scaling exponent, γ) indicated a weak dependence of t_1/2_ on initial monomer concentration (Fig. 2A and 2B insets). This observation indicates fibril breaking being the prominent mechanism for the generation of new seeds. A corresponding time-dependent structural changes in the α-Syn secondary structure were monitored using far-UV CD spectroscopy (Fig. 2C and 2D). The CD spectral peak of native α-Syn was negatively centered around 196, confirming its predominantly random conformation. In MP, upon incubation, the 196 nm peak of α-Syn gradually reduced in intensity, and a new small negative peak emerged around 220 nm, indicating the formation of secondary structures (Fig. 2C).

**Fig. 1:**
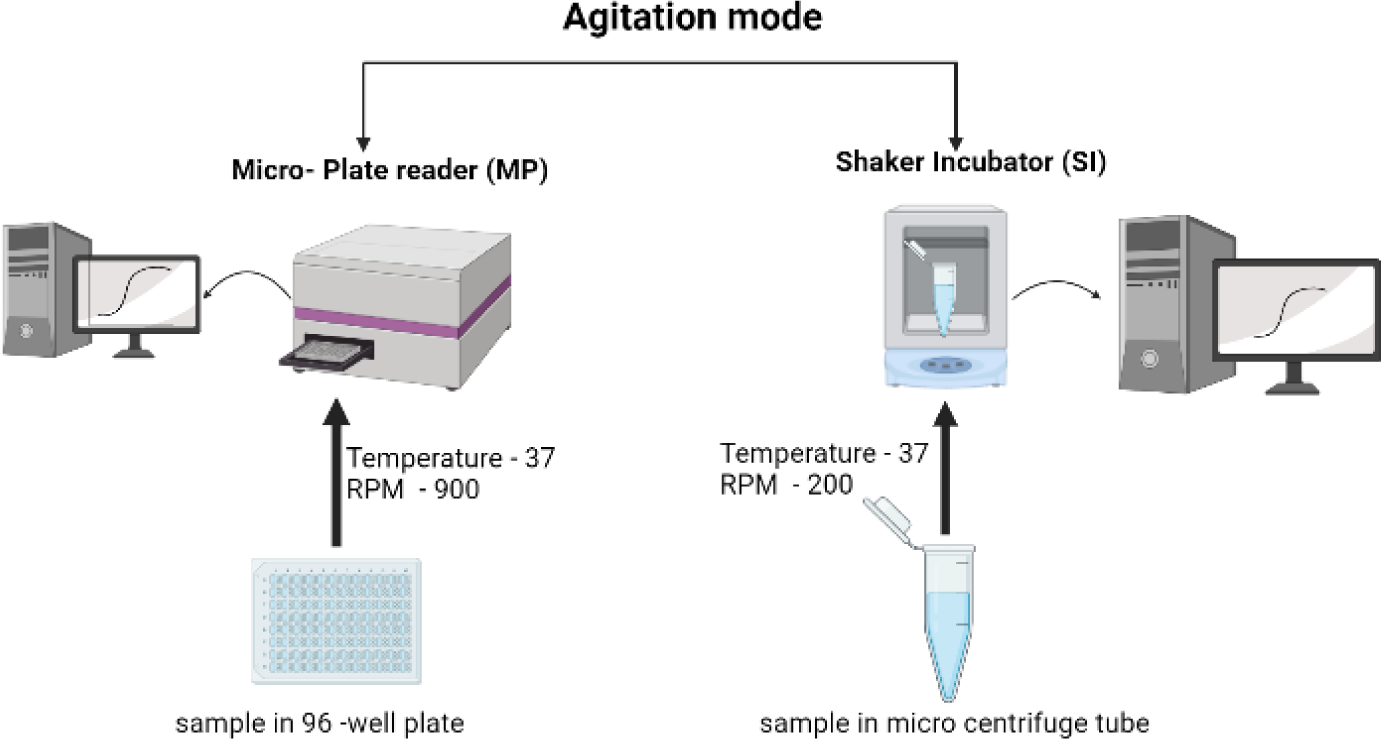
Two agitation modes used for to study the in vitro aggregation of α-Syn in this work.

**Fig. 2:**
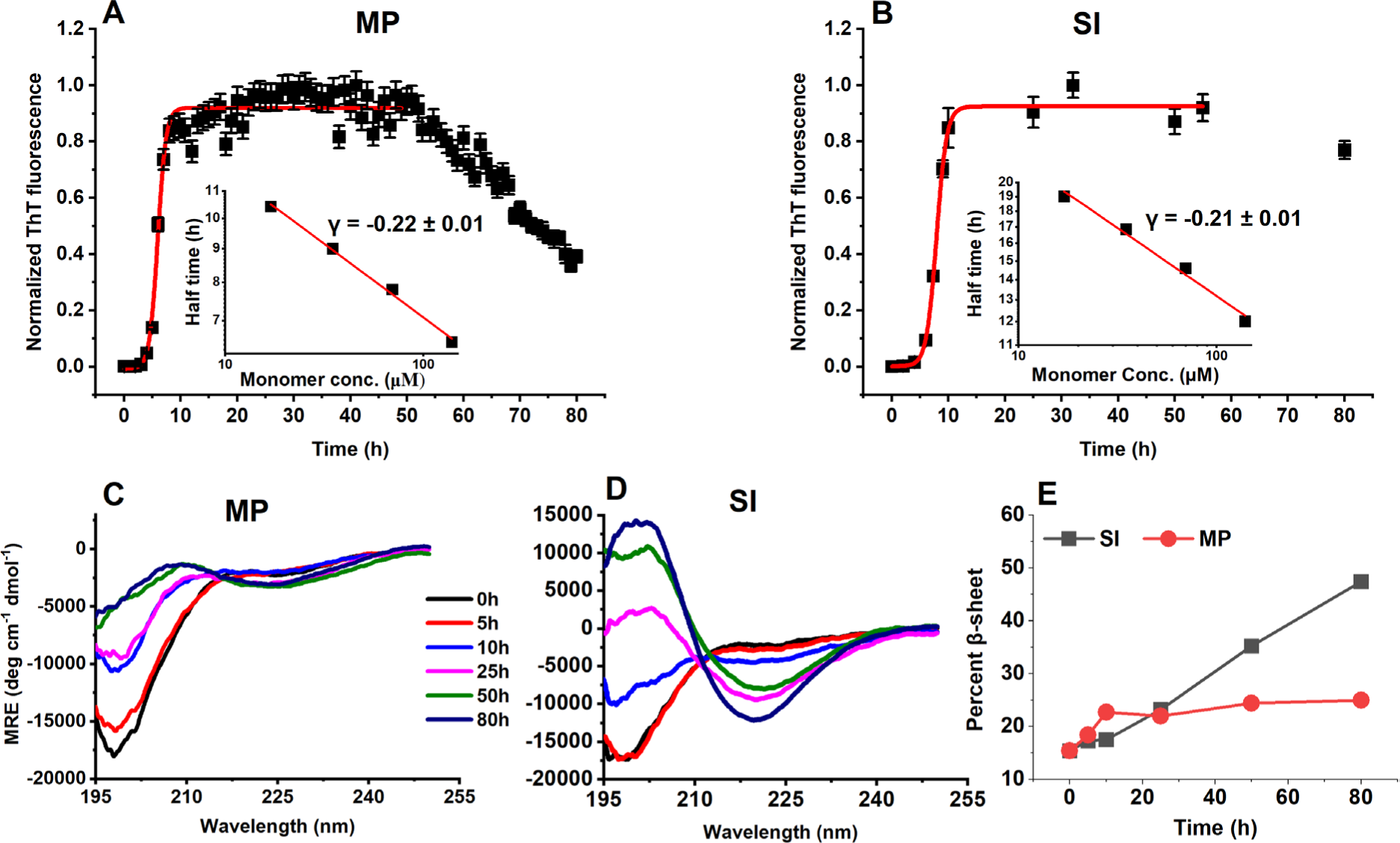
Aggregation kinetics of α-Syn. ThT fibrillation kinetics of 1 mg ml^−1^ α-Syn (70 μM) was carried out separately in a 96-well microplate reader (MP) and a shaker incubator (SI) in 10 mM Na-phosphate buffer, 100 mM NaCl, 20 μM ThT, and 0.02% sodium azide at 37°C. The half time of the aggregation kinetics (t1/2) was plotted against different monomer concentration using double logarithmic scale (panel A and B) **(A)** The protein sample was agitated at 900 rpm in MP, and **(B)** 200 rpm in SI. The red line represents sigmoidal fit. Error bars: mean ±SEM of n = 4 samples. **(C)** and **(D)** A time-dependent change in the secondary structure was monitored using far-UV CD for MP and SI, respectively. **(E)** The time evolution of β-sheet during aggregation demonstrates the random coil to β-sheet transition during aggregation.

**Table 1:** Kinetic parameters of MP and SI aggregation derived by fitting the raw data in the sigmoidal model.

| | amplitude | lag time (h) | $t_{1/2}$ (h) |
| --- | --- | --- | --- |
| **MP aggregates** | $0.92 \pm 0.04$ | $4.52 \pm 0.63$ | $5.98 \pm 0.16$ |
| **SI aggregates** | $0.92 \pm 0.03$ | $6.13 \pm 0.39$ | $7.93 \pm 0.19$ |

On the contrary, the SI incubated sample, showed a gradual reduction in 196 nm peak from 10 h onwards, with a concomitant emergence of positive and negative peaks around 196 and 218, respectively, indicating the formation of β-sheet structures (Fig. 2D). The structural transition occurred until 50 h, followed by a slight structural change upon further incubation till 80 h in the case of SI sample. At 80 h, while the CD structure of MP incubated sample remained essentially random, that of SI incubated sample was prominently β-sheet rich. Further, as a measure of β-sheet content in MP and SI samples, we deconvoluted CD spectra using CONTIN software and plotted percent β-sheet content with respect to time (Fig. 2E). The β-sheet content of the MP sample remained largely unchanged from 5 h onwards, whereas that of SI sample increased with time. From ThT and CD data, it was interesting to note that: 1) The CD spectra of MP aggregates remained mostly random, much beyond its ThT saturation point (12 h); 2) The secondary structural evolution of α-Syn in SI continued to occur beyond its ThT saturation point (10 h) upto 80 h.

### 2.2. The end-stage aggregates in two agitating modes were polymorphic

We characterized the nanoscale morphology of aggregated α-Syn at two time points that lie beyond ThT kinetics saturation points: 1) 50 h when most of the CD structure change had taken place in both MP and SI conditions; and 2) 80 h, an extended time point, beyond which no further structural change take place in the case of SI aggregates. At 50 h, MP aggregates were in the form of 0.5-1.5 μm length irregular clumps, from which 3-4 nm thick fibrils were seen emerging (Fig. 3A). Upon further incubation, upto 80 h, there was a slight change in the morphology of MP incubated sample, with a few clumps dissolve into small fibrillar species which were still attached together. On the other hand, SI incubated samples displayed straight morphology with fibril length in the range of 5-10 μm, with an average thickness of 4 nm. Upon further incubation, upto 80 h, we did not observe any appreciable change in the morphology and average dimensions of the SI fibrils.

**Fig. 3:**
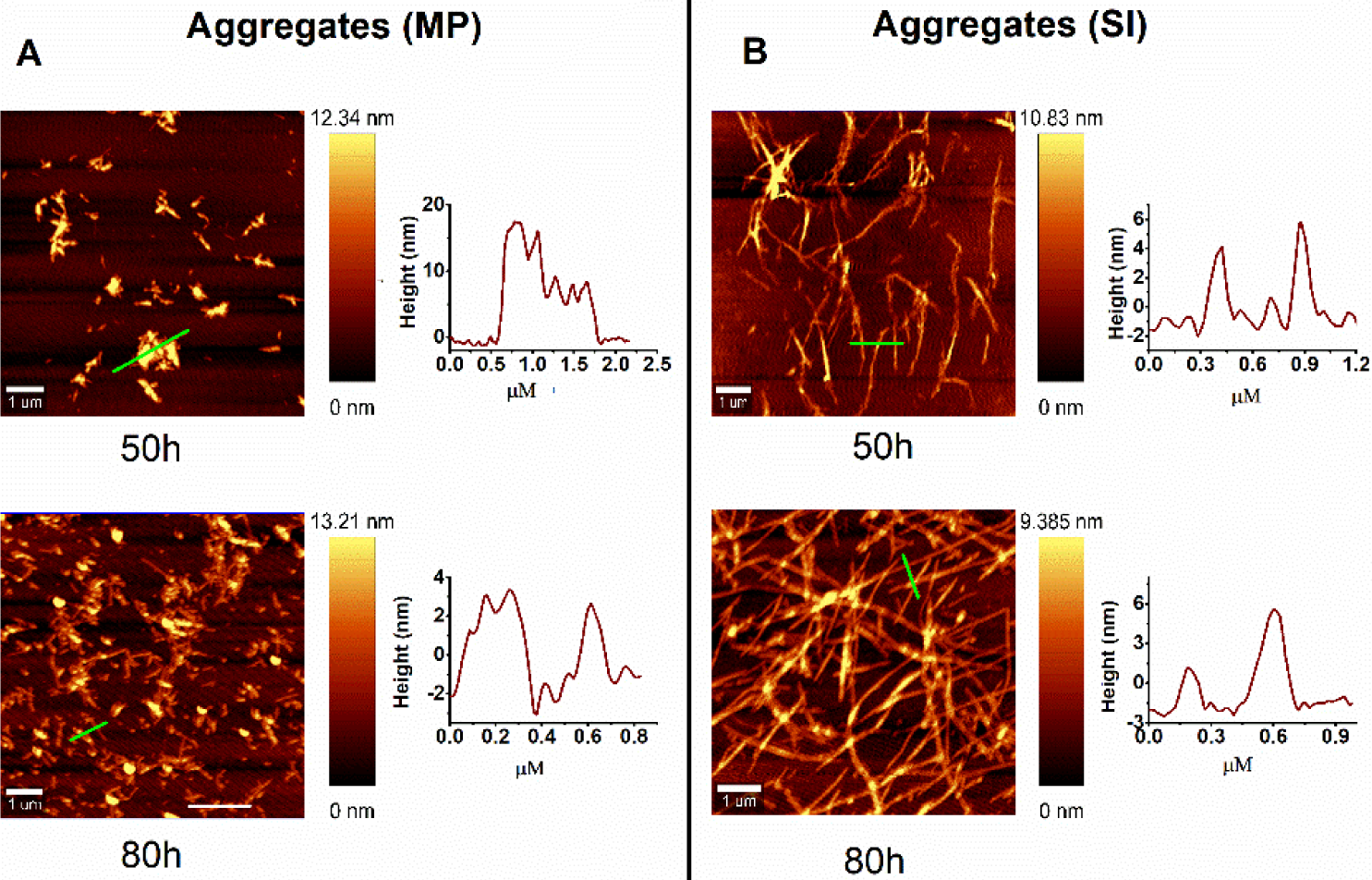
AFM images of MP and SI incubated α-Syn. Morphology of MP **(A)**, and SI **(B)** incubated α-Syn end-stage aggregates at 50 and 80 h, along with their respective height profiles shown alongside each image. The height profile was calculated by obtaining a line profile (green line). Scale bar: 1 μm.

### 2.3. Vibrational spectroscopies confirm underlying structural differences among two polymorphs

Vibrations in the molecular bonds in proteins are characterized by a specific vibrational frequency and bond polarizability. Both frequencies and polarizability are sensitive to the bond environment, which dependents on the protein fold [34, 35]. A change in these physical quantities is well exploited in vibrational spectroscopies. Raman scattering and infrared (IR) absorption are the two well-established vibrational spectroscopic tools that can unravel differences in the arrangements of polypeptide chains in different aggregate polymorphs [36, 37]. These two methods complement each other in deciphering different parameters of bond vibrations and, when used in conjunction, can give rise to important insight into the structural signatures of the molecule under study. Each aggregate polymorph gives rise to a characteristic Raman scattering or IR absorption signatures which serve as its fingerprint. These techniques are considered complementary to CD spectroscopy in case where solution is non-homogenous due to aggregation.

Vibrational Raman scattering arises primarily from C=O stretching and in-plane N-H bending in amide I region (1600 – 1700 cm^−1^); and C-C, C-N bond vibration, and C=O in plane bend in the amide III region (1200-1400 cm^−1^) [38, 39]. The Raman spectra for both MP and SI polymorphs of α-Syn were collected in a wave number range encompassing both amide I and amide III regions (Fig. 4A). The amide I region of monomeric α-Syn consisted of a major peak centred around 1667 cm^−1^, which was slightly shifted to 1666 cm^−1^ in MP and remained the same for SI (Fig. 4B). However, a narrowing of the peak was visible upon aggregation. The Raman shift corresponding to this wavelength range is assigned to β-sheets. The area under the curve for these β-bands increased slightly for aggregated samples, with maximum values for SI incubated samples.

**Fig. 4:**
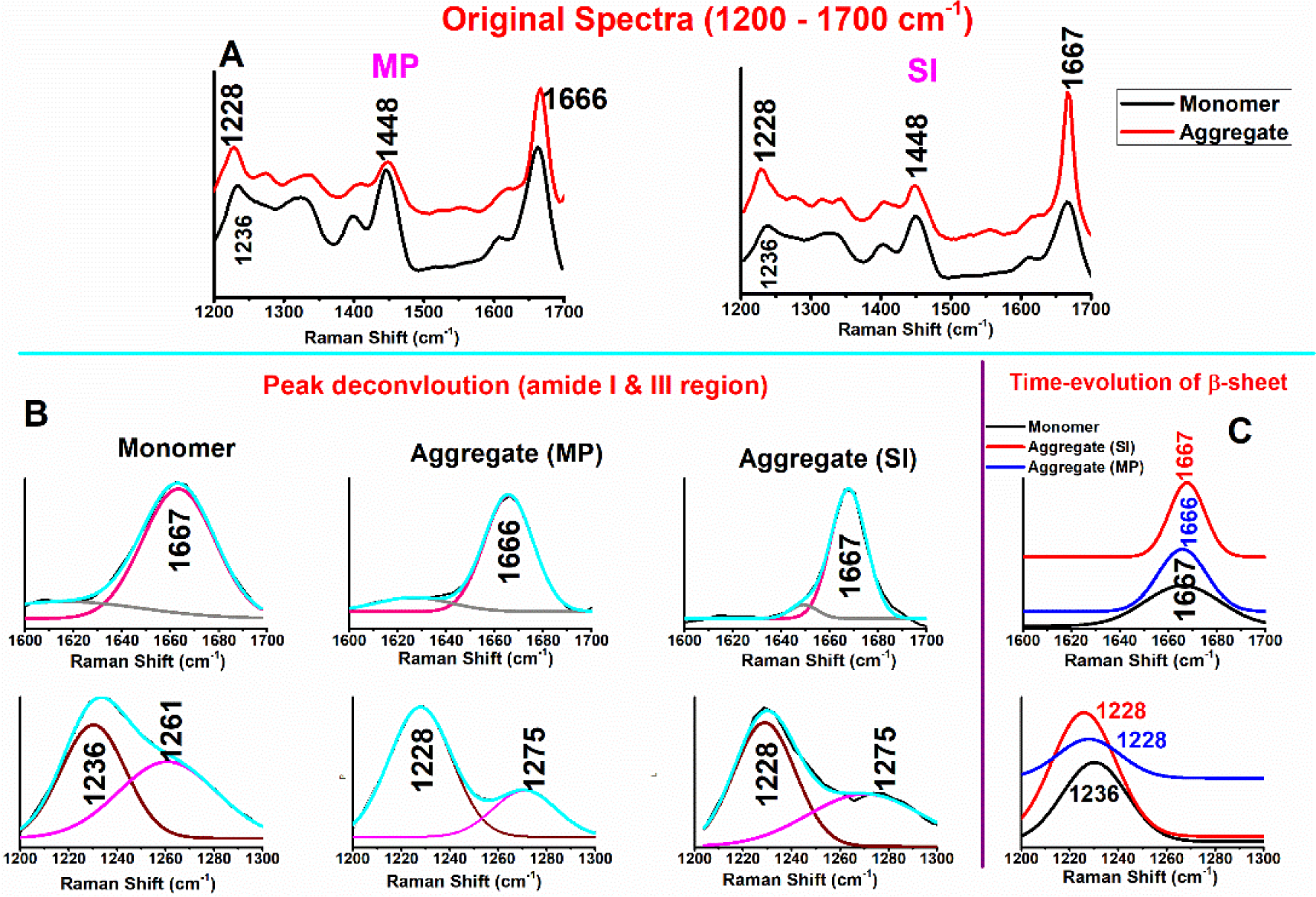
Raman scattering of α-Syn aggregates recorded on a confocal Raman imaging system. **(A)** Complete Raman spectra of α-Syn aggregates formed in MP and SI post 50 h incubation. A 0 time point monomer spectra is included as the control. **(B)** Amide I (upper panel) and amide III (lower panel) regions of aggregates are compared with monomer. **(C)** Band regions corresponding to β-sheet in the amide I (∼1666 cm^−1^) and amide III (1228-1236 cm^−1^) for the end-stage aggregates are shown for comparison. The sharpening of bands (reduction of FWHM values, table..) indicates the formation of β-sheet [40, 41].

We also derived the full width at half maximum (FWHM) values of each Raman band, which indicates the sharpness of the peak. FWHM is the difference between two X values (wavelength) on the curve at which the Y value (intensity) is half the maximum. Previous reports have highlighted that upon aggregation of α-Syn, the peak corresponding to 1666 cm^−1^ narrows, as quantified through a reduction of FWHM values (Fig. 4B, Table 2) [40, 41]. The FWHM value of unaggregated control was 35.17 cm^−1^, much broader than the aggregated samples. Among two aggregated polymorphs, the FWHM value for the 1666 cm^−1^ band for SI aggregate was sharper (17.3 cm^−1^) than that of MP aggregates (23.6 cm^−1^), confirming a more structured conformation of SI aggregates (Fig. 4, Table 2). A minor peak corresponding to 1616 cm^−1^ in the monomer was also observed that might be contributed by the tyrosine residues [42]. This peak was shifted to 1626 and 1649 cm^−1^ for MP and SI aggregates, respectively, corresponding to α-helix.

**Table 2:** Raman scattering peak deconvolution data reveal two prominent bands each in the amide I and III regions. The narrowing of different bands denotes the formation of cross-β structure and is numerically represented by the full width at half maximum (FWHM).

|  | monomer |  |  | MP aggregate |  |  | SI aggregate |  |  |
| --- | --- | --- | --- | --- | --- | --- | --- | --- | --- |
| | Raman shift ($\text{cm}^{-1}$) | Peak area (%) | FWHM ($\text{cm}^{-1}$) | Raman shift ($\text{cm}^{-1}$) | Peak area (%) | FWHM ($\text{cm}^{-1}$) | Raman shift ($\text{cm}^{-1}$) | Peak Area (%) | FWHM ($\text{cm}^{-1}$) |
| **Amide-I (1600-1700)** |  |  |  |  |  |  |  |  |  |
| Peak 1 | 1663 | 78.3 | 35.17 | 1666 | 84.1 | 23.6 | 1667 | 92.3 | 17.3 |
| Peak2 | 1616 | 21.6 | 74.9 | 1626 | 15.8 | 36.9 | 1649 | 7.6 | 13.6 |
| **Amide – III (1200-1300)** |  |  |  |  |  |  |  |  |  |
| Peak 1 | 1236 | 49 | 31.2 | 1228 | 73.3 | 30.4 | 1228 | 55.9 | 29 |
| Peak 2 | 1261 | 50.8 | 47.8 | 1275 | 26.5 | 30.7 | 1275 | 44 | 55 |

The Raman shift corresponding to the amide III region also provides an approximation for the secondary structure content. The 1236 cm^−1^ band in the monomer shifted to 1228 cm^−1^ during aggregation, hinting at the formation of β-sheets. We also observed a broad peak centred at 1261 cm^−1^ and 1275 cm^−1^ for the monomer and aggregates, respectively. However, a high FWHM of these peaks precluded their assignment to a particular structural motif.

To further confirm the structural underpinning in the structural differences among the two aggregates, we conducted an attenuated total reflection Fourier transform infrared spectroscopy (ATR-FTIR) analysis of different conformations of α-Syn. The FTIR signature is produced by the absorption of radiations by the bonds in protein molecules in the mid-infrared region. It is sensitive towards detection of polarized bonds (functional groups with strong dipoles) [43, 44]. Examples include heteronuclear groups such as C=O, N=O, and O-H. The H_2_O interferes strongly with the spectral acquisition in the amide I region (1600-1700 cm^−1^), where protein produces its characteristic signature [45]. To mitigate this issue, we first pelleted the aggregated protein through centrifugation and washed the pellet in D_2_O to get off residual H_2_O. We then carried out an ATR-FTIR analysis of the aggregates. ATR mode enabled us to analyze the insoluble/solid aggregated samples, which were otherwise not suitable for study through transmission mode FTIR, which requires the sample to be in a solution state.

The spectral deconvolution of FTIR spectra of α-Syn monomer reveals a prominent peak centred around 1643 cm^−1^ that corresponds to disordered conformation (Fig. 5A, Table 3). A minor peak with centre at 1682 cm^−1^ was also observed alongside, which indicated the presence of β-turns. The deconvoluted spectrum of MP aggregate revealed three peaks centred at positions 1666, 1644, and 1624 cm^−1^, corresponding to turns, random coil, and β-sheets, respectively (Fig. 5B, Table 3). In comparison, the corresponding peaks in the SI aggregates were 1662, 1647, and 1625 cm^−1^ (Fig. 5C, Table 3). A particular difference of FTIR signals between MP and SI aggregates FTIR was the broadening of 1662 cm^−1^ peak (corresponding to turns) and reduction of the peak corresponding to the random coil (1647 cm^−1^). From FTIR spectral analysis, a significant random coil structure was evident in MP aggregates that was negligible in SI aggregates.

**Fig. 5:**
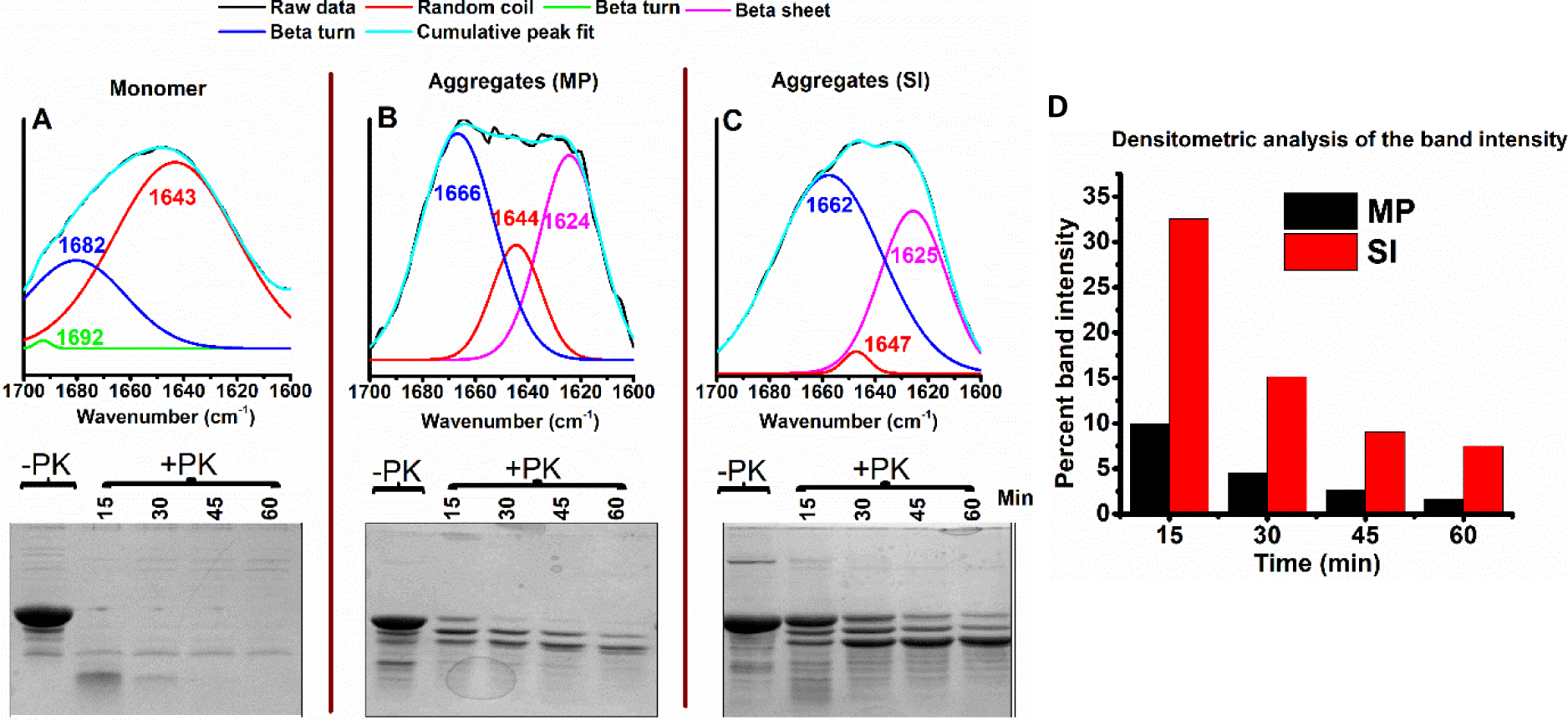
Comparing amyloid cores of end-stage aggregates. **(A), (B), and (C)**. Upper panel, FITR; lower panel, PK digestion profiles of monomers and aggregates of α-Syn. Black curves are experimental signals; Cyan curves are fits. Data were fitted using a Gaussian model. Signals under the curve represent deconvoluted experimental curve-fit signals bands used to determine structural content in a given sample. The purple, blue and red lines represent the contribution of β sheet structures, β turn, and - random coil, respectively. α-Syn becomes PK-resistant upon aggregation in vitro. α-syn monomer and its MP and SI aggregates were incubated with PK for 1h. Treated samples were taken out at different time intervals and mixed with SDS-Laemmli’s buffer to stop the reaction. Tricine-SDS (16%) was used to visualize the band profiles of PK digested samples. The PK digestion of aggregates reveals three prominent bands. **(D)**. Densitometric quantification of band 1 of MP and SI aggergates reveal that for SI samsple, it is significantly PK stable.

**Table 3:** FTIR peak deconvolution to delineate the relative percentages of different secondary structural motifs in monomer and aggregates.

|  |  | **Area under the peak (%)** | **Structural assignment** |
| --- | --- | --- | --- |
| **monomer** | Peak 1 (1643) | 79 | Random |
| | Peak 2 (1692) | 0.11 | $\beta$ - turn |
| | Peak 3 (1682) | 21 | $\beta$ - turn |
| **microplate (MP) aggregates** | Peak 1 (1624) | 35.6 | $\beta$ - sheet |
|  | Peak 2 (1644) | 16.3 | Random |
| | Peak 3 (1666) | 48 | $\beta$ - turn |
| **shaker-incubator (SI) aggregates** | Peak 1 (1625) | 33.4 | $\beta$ - sheet |
|  | Peak 2 (1647) | 1.67 | Random |
| | Peak 3 (1662) | 64.9 | $\beta$ - turn |

To understand the sensitivity of the two polymorphs towards proteolysis, we carried out time-dependent protease digestion of both the aggregates through proteinase K (PK) and compared the digestion pattern with that of monomer (Fig 5, lower panel). PK is a serine endoprotease with a low amino acid specificity and can hydrolyze any exposed polypeptide segment that lies outside the rigid protein amyloid core [46, 47]. It is therefore traditionally used to identify structural polymorphs of amyloids, as different polymorphs reveal different signature protease-resistant cores upon PK digestion. A time-dependent PK digestion profile of monomers revealed that it was almost degraded beyond 30 min. The PK digestion profiles of MP and SI aggregates resulted in three major bands at 15 min time points, with the topmost band corresponding to the full-length α-Syn. For the subsequent time point, beyond 30 min, the band corresponding to full-length protein was degraded in MP aggregate, while it remained resistant to proteolysis in SI aggregates (Fig. 5D). It is noteworthy that the total quantity of two aggregated polymorphs in the gel was a little different (for untreated sample), despite that same volume of aggregated samples were analyzed. These samples were mixed with an identical volume of Laemmli buffer, boiled for 5 minutes, and centrifuged at 10,000 g for 5 minutes.

From there, 10 μl supernatant was loaded onto the gel. It might be possible that after centrifugation, a differential amount of sample remained in the supernatant. Along with FTIR results, PK analysis confirms the presence of a loosely packed structural motif in MP aggregates.

### 2.4. MP aggregates were in kinetically trapped conformation that can undergo further structural rearrangement

The MP aggregates display unusual CD spectra with significant random coil structures and small clumped aggregate morphology (Fig. 2C & 3A). On the contrary, SI aggregates show a typical β-sheet CD signature and fibrillar morphology characteristic of amyloid fibrils (Fig. 2D & 3B). Time-dependent CD analysis (Fig. 2D) reveals that in the case of SI aggregates, a similar random coil-rich CD structure was observed at 10 h when the log phase of its aggregation kinetics commences. To check whether MP aggregates were in structurally trapped conformation, in the first experiment, we added preformed SI aggregate seeds in α-Syn monomers at 5, and 10% monomer equivalent concentrations, and carried out the aggregation reaction in microplate, and analyzed the end-stage aggregates through far UV CD spectroscopy (Fig. 6A). The CD of aggregates that were seeded with preformed SI aggregates showed a typical β-rich structure, compared to that of unseeded (de novo) reaction that showed a mix of random coil and β-sheet structure.

**Fig. 6:**
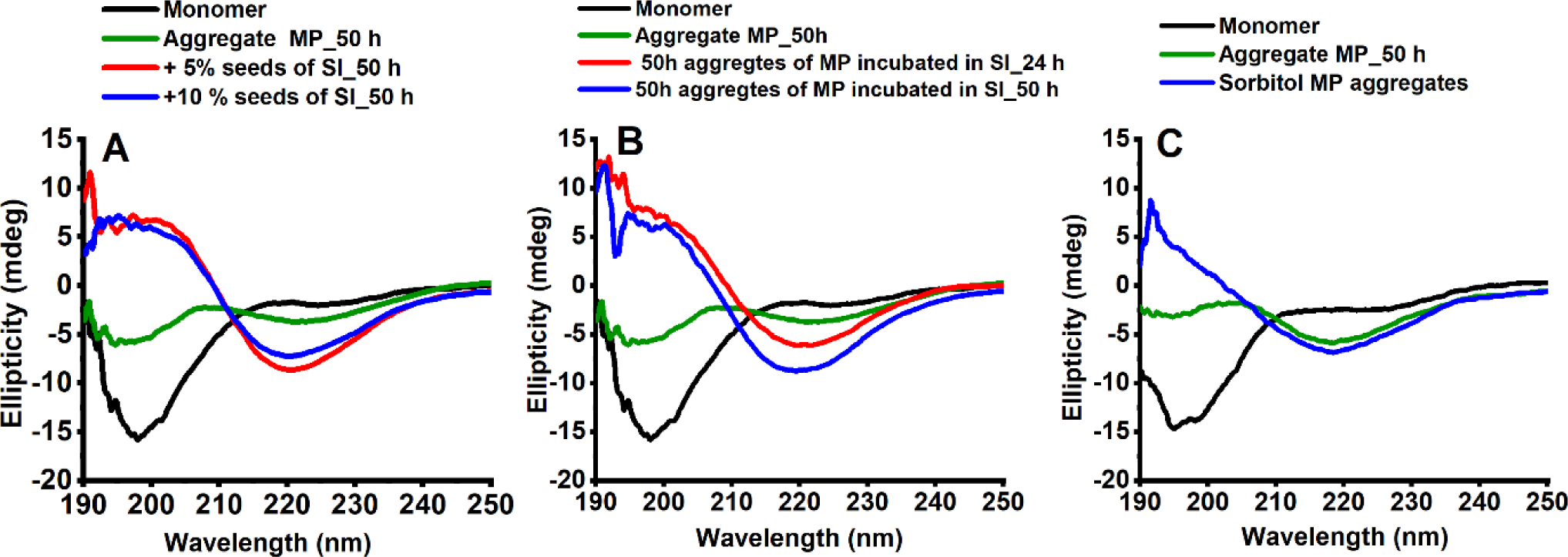
Changes in aggregation conditions alters aggregation landscape of MP aggregates. **(A)** Initiation of aggregation reaction of α-Syn in MP in the presence of preformed β-sheet rich SI seeds at 5% and 10% monomer equivalent concentration results in the formation of β-sheet rich aggregates. **(B)** Transfer of end-stage MP aggregates having random coil + β-sheet conformation to SI conditions results in their structural conversion to β-sheet rich aggregates. **(C)** In the presence of 1.5 M sorbitol, an osmolyte, α-Syn forms β-sheet-rich aggregates in MP conditions.

In a subsequent experiment, we checked whether end-stage random coil/β-sheet rich MP aggregates were in structurally trapped conformation. We subjected the end-stage MP aggregate to agitating conditions in a shaker incubator (SI), followed by their analysis through CD after 24h and 48h post-SI incubation (Fig. 6B). We found that within 24 h, a further structural conversion occurred as discernible through the acquisition of β-sheet CD signature, like that of aggregates formed in SI conditions. There was a little structural change upon further incubation till 50 h. Expectedly, the CD spectra of the MP control (96 h incubated sample in MP) remained in random/β conformation.

Next, we carried out the aggregation α-Syn monomers in a microplate, in the presence of 1.5 M sorbitol, a cellular osmolyte (Fig. 6C). Osmolytes are small organic molecules that are preferentially excluded from the polypeptide backbone and favour a compact structure [48, 49]. This property of osmolytes imparts stability to the globular proteins that prefer to remain in compact folded states [50]. For intrinsically disordered proteins (IDPs), however, the addition of osmolytes results in the compaction of the natively disordered chain, an unfavourable conformation for an IDP [51]. Such a relatively compact structure of IDPs may result in intermolecular association. Alternatively, IDPs may also undergo intermolecular association to reduce the chain exposure to the osmolytes. Either way, in general, the effect of osmolytes on IDPs may also result in their oligomerization or aggregation, as evident in earlier reports on α-Syn and other IDPs [52–54]. We observed that the end-stage aggregates formed in the presence of 1.5 M sorbitol showed a β-sheet signature compared to control aggregates, which showed a mix of random coil and β-sheet.

Finally, to understand the kinetics of MP and SI-seeded aggregation of α-Syn in MP-shaking conditions, the quantitative parameters of three aggregation processes were compared (Fig. 7 and Table 4). Both MP- and SI-aggregates were able to seed the fresh reaction of α-Syn in the microplate, as evident from the reduction of lag time upon seeding (Table 4). To compare the maximum rate of the aggregation reactions of three processes, we derived the slope of the kinetic curves at t_1/2_ where 50% of the aggregation takes place. The maximum rate (at t_1/2_) of unseeded and MP-seeded reactions were comparable implying that only the lag time of the reaction was reduced upon seeding and overall aggregation pathway remained the same. We also calculated the initial reaction rate by deriving the slope at t_0_. We found that the initial rate of SI-seeded reaction was significantly higher than that of MP-seeded reaction, indicating faster recruitment of monomers to the SI-aggregates. However, the maximum rate of SI-seeded reaction was less, when compared to unseeded and MP-seeded reactions pointing to a possible modulation of the aggregation pathway upon SI-seeding.

**Fig. 7:**
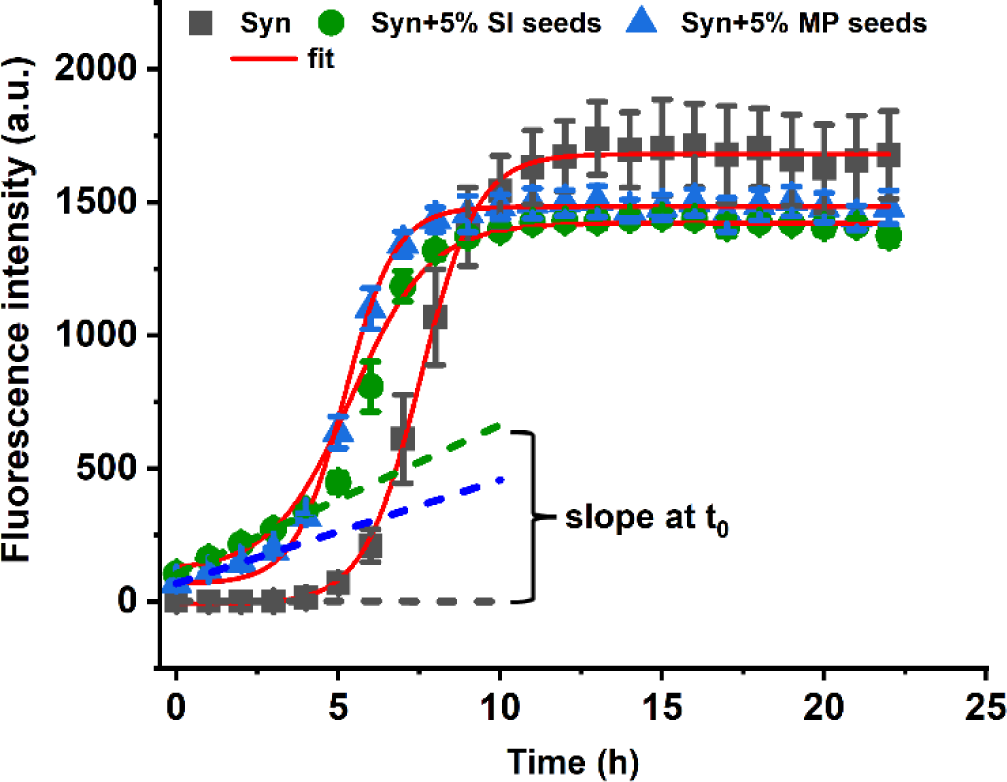
Aggregation kinetics of α-Syn in MP conditions in the presence of MP- and SI-derived seeds. Seeding was performed by adding 5% molar equivalent of end-stage aggregates in fresh monomers of α-Syn and the reaction mixture was agitated in microplate conditions. The initial slope of the kinetics at 0 time (t0) is indicated. Error bars: mean ± SEM of n = 4.

**Table 4:** Kinetic parameters of α-Syn aggregation seeded with MP- and SI-derived seeds.

| | Lag time | Initial slope of reaction at $t_0$ (ThT fluorescence/h) | Maximum slope of reaction (ThT fluorescence/h) |
| --- | --- | --- | --- |
| **Control (no seed)** | $5.14 \pm 0.32$ | $-0.17 \pm 0.23$ | $458 \pm 17$ |
| **MP-seed** | $3.66 \pm 0.16$ | $38.9 \pm 1.5$ | $454 \pm 26$ |
| **SI-seed** | $3.02 \pm 0.23$ | $56 \pm 0.33$ | $268 \pm 9$ |

With these observations, we conclude that the microplate shaking conditions promote the formation of end-stage aggregates in a kinetically trapped state, wherein a significant portion of the α-Syn chain remains in disordered conformation. This kinetic trap is bypassed by adding SI seeds and sorbitol. Alternatively, when subjected to SI conditions, the preformed MP aggregates can overcome the kinetic barrier and undergo a further structural rearrangement.

## 3. Discussion

In this study, we seek to understand in vitro aggregation mechanism of α-synuclein through two complementary tools. We employed the two most used agitation modes in our research that are being extensively utilized to study *in vitro* amyloidogenesis of different proteins. We used time-dependent ThT fluorescence as a direct indicator of cross-β motif formation-a hallmark of amyloidogenesis of proteins. We correlated ThT data with far-UV CD spectra that report a global change in the secondary structure accompanying amyloidogenesis of α-synuclein. In general, enhancement in ThT fluorescence during aggregation correlates with the acquisition of cross-β structure. This model is essentially valid for nucleation-dependent polymerization, wherein the formation of a β-sheet nucleus precedes the subsequent recruitment of native monomers to the growing fibril end followed by conversion of the native structure of the incoming monomers to β-sheet [55–57]. Spectroscopic manifestation of such a model is the concomitant increase in β-sheet content (monomer structure conversion upon assembly) and enhancement in ThT fluorescence (formation of a cross-β motif with the nucleus). The saturation phase in ThT kinetics marks the depletion of soluble monomers and an end to the aggregation process. This concordance of ThT fluorescence and CD-monitored structural conversion may not always be valid for the following reasons. As per the widely posited mechanism, ThT is a class of molecular rotor whose rotation is confined upon binding to the cross-β motifs of amyloids [58]. Such rotational confinement preserves its excited state, which results in a high quantum yield of its fluorescence. The availability of the number of binding sites for ThT on an aggregate or its precursor state may vary depending on their conformation. Such a difference in the ThT binding sites is also apparent from different ThT binding of various polymorphs of α-Syn [59]. A recent study highlights the presence of ThT negative α-Syn fibrils [60]. In our case, an initial increase in ThT fluorescence may correspond to the acquisition of β-sheets by α-Syn monomers. Upon further incubation, two processes might happen simultaneously: 1) generation of ThT binding sites by the formation of cross β-sheets among remaining monomers and prefibrillar species (oligomers), and 2) sequestration of ThT binding sites by the assembly of oligomers into fibrils. So, while CD readouts at 218 nm for the bulk α-Syn in SI mode increased with time, that of ThT became constant. For the proteins that follow nucleation-conversion-polymerization model, monomers first collapse in a native-like or disordered assemblies, followed by their further structural rearrangement into β-sheet rich aggregates [24–27, 61]. For α-Syn, both the aggregation models are proposed [12, 22–24]. In our observation, end-stage MP-derived aggregates showed random CD spectra (Fig. 2C) and irregular AFM structures (Fig. 3A). Owing to irregular clump-like morphologies, these aggregates did not resemble oligomers or fibrils, which have a defined shape and thickness. Further, unlike early oligomers and end-stage fibrils, which show a well-defined β-sheet content, these aggregates showed a largely random CD structure. This observation proved that they were in structurally collapsed form, in which monomers did not acquire a perfect β-sheet conformations. The acquisition of β-sheet CD structure upon shifting these aggregates in SI condition revealed their kinetically trapped conformation. This observation hinted that in the case of MP, α-Syn aggregation was not through a simple nucleation dependent polymerization.

During protein folding, the conformational search is biased towards a single energy minimum in a funnel-like energy landscape [62]. This single energy minimum corresponds to a folded functional state of that protein, which is robust to a minor change in solution conditions such as salt concentration, pH, temperature, etc. For protein amyloidogenesis, however, simulation experiments have shown that the conformational landscape is not biased towards a single state and appears like a flat rough landscape with numerous minima [63]. According to another view, a combination of the funnel and flat rugged landscape exists for the amyloid landscape [64]. The rough and unbiased landscape of amyloidogenesis is evident from the co-existence of different assembly products such as amyloid fibrils, and microcrystals in the same aggregate preparation [65, 66]. However, proof of the combination of the funnel (biased) and unbiased rough landscape comes from a study where low hydrodynamic solution stress generated heterogenous α-Syn fibrils (unbiased rough component). In contrast, in the same study, high-stress conditions resulted in the fibrils of uniform morphology (funnel component) [67]. On biased and unbiased energy landscapes, the local energy minima act like kinetic traps, separated from other more prominent minima by a higher energy barrier. In other words, the most frequent fibril polymorphs are not necessarily the most stable [68–70]. For these reasons, kinetic control is hypothesized to play a significant role in amyloid polymorphism.

A fibril polymorph may not be the final state and changing its physico-chemical environment can help it overcome the kinetic barrier to adopt an alternate conformation. In recent work, such an observation was reported for in vitro generated lysozyme fibrils, wherein two polymorphs were interconvertible with a change in hydrophobicity of the buffer solution [71]. Polymorph interconversion is usually a rare phenomenon because for such interconversion to occur, the stable, rigid amyloid core must disassemble before reassembling, which is a kinetically unfeasible process. However, the flexible regions outside the rigid core can modulate global change in fibril characteristics such as toxicity and can readily undergo structural change. For example, in the case of polyQ fibrils formed in Huntingtin’s disease, the C-terminal proline-rich domain undergoes various degrees of dynamic entanglement to give rise to two different polymorphs of fibrils [72]. Notably, the polyQ fibril core remains unchanged in two polymorphs. In our study, too, we opine that MP and SI aggregates were not different per se, *i.e.*, their structural core was not diverse, as evident from the following observations. 1) The three FTIR bands of both aggregates were at a similar position and differed only in the area covered under each band (Fig. 5; Table 1), 2) The proteinase K digestion experiment revealed three prominent bands on SDS-PAGE, in both aggregates, albeit with different intensities. Instead, they were kinetically separated because the end-stage random-coil MP aggregates were converted into β-sheet aggregates upon subjecting to SI agitating conditions (Fig. 6B).

The kinetic barrier for α-Syn aggregation is relatively high, as it requires vigorous shaking for its in vitro aggregation and does not aggregate for days if not agitated. This observation contrasts with other aggregation-prone polypeptides such as TDP-43, yeast prion protein, Aβ42, etc., which undergo spontaneous aggregation even under quiescent conditions [73–75]. It is hypothesized that the positive and negative N- and C-terminal domains of α-Syn interact to protect the aggregation-prone central NAC domain [11]. It is noteworthy that the high kinetic barrier for α-Syn aggregation is not just limited to the nucleation stage. Nucleation is a rate-limiting step that involves a slow association of monomers in an aggregation-competent high-energy oligomeric nucleus, which is a thermodynamically unfavorable process [56]. If nucleation were the only rate limiting factor, then upon nucleus formation, a further elongation step would have been spontaneous, and only fibrils should have been observed consistently. However, different on-pathway or off-pathway metastable prefibrillar species of a varying hierarchy of α-Syn are abundantly reported and theorized to be in kinetically trapped states. These include oligomers [76, 77], annular protofilaments [78, 79], and long protofilaments [80]. Such prefibrillar species are proposed to be more toxic than mature fibrils.

The crowded molecular environment of the cell can potentially increase or decrease such kinetic barriers by mechanisms specific to the crowding molecules. For example, macromolecular crowder vs small osmolytes, lipid vs protein, charged vs uncharged molecules, etc. In one study, lipid-induced kinetically trapped protofibrils were observed [80]. While in another work, Ficoll70, a macromolecular crowder that provides excluded volume effect, increased the kinetic rate of α-Syn aggregation [81]. In contrast, its monomeric building unit, sucrose, an osmolyte, slowed down the kinetics. In our case, however, sorbitol, a small molecule polyhydric alcohol osmolyte, helped α-Syn overcome its kinetic barrier in microplate conditions. Osmolytes are preferentially excluded from the protein backbones and thus favor a compact conformation of different proteins [49, 82]. For intrinsically disordered proteins, which are thermodynamically unfolded, the addition of osmolytes can either induce a partially folded structure (intramolecular interactions) or induce oligomerization (intermolecular interactions) [10, 52]. We rule out the former case (Fig. S4) and opine that the addition of sorbitol in our study might reduce the kinetic barrier and promote intermolecular interactions (Fig. 8).

**Fig. 8:**
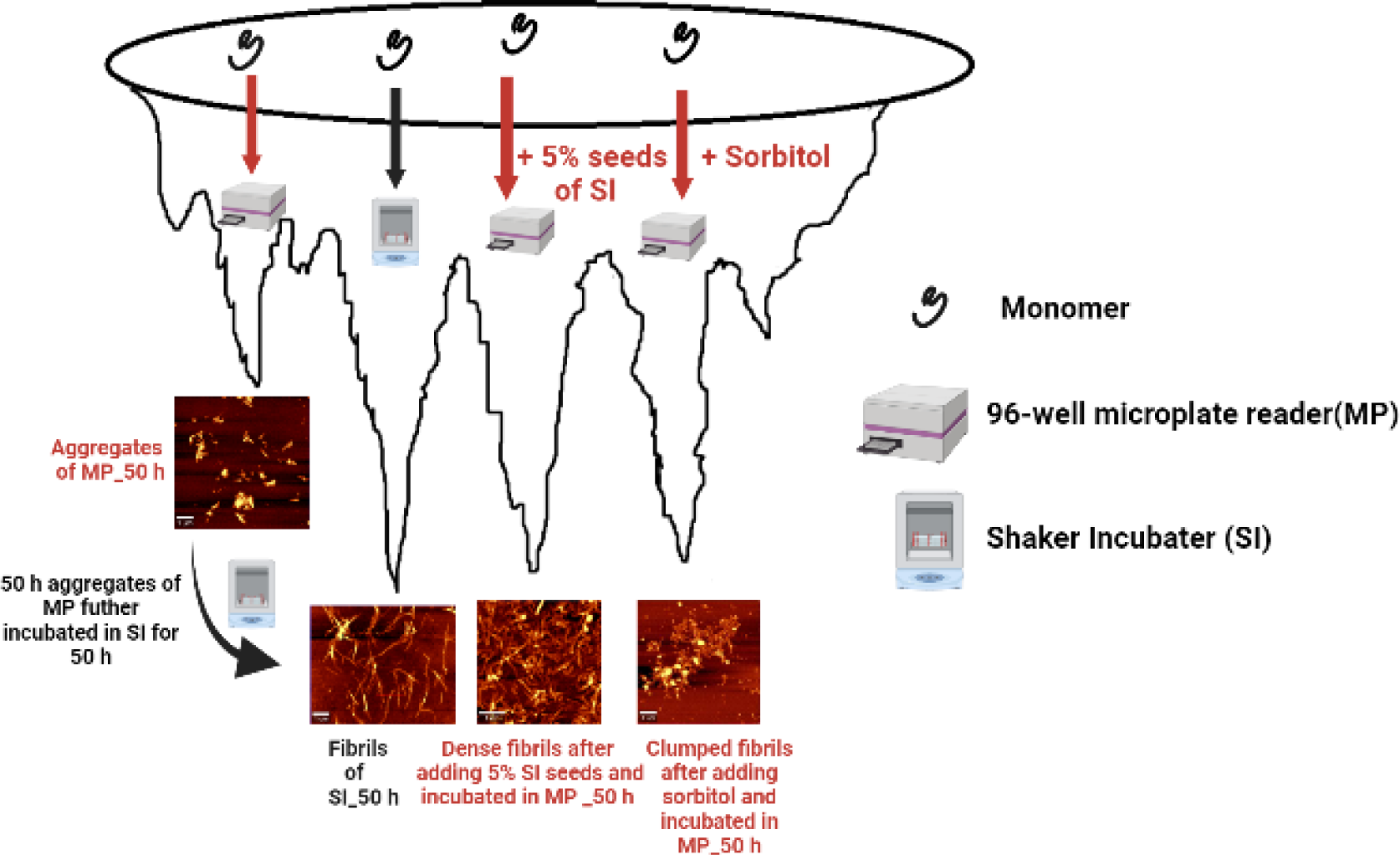
Schematic representation showing the kinetic partitioning of α-Syn aggregates formed in different agitation modes, and the crossing of such barrier leading to their interconversion.

In summary, we generated kinetically trapped amyloid aggregates of α-Syn that showed a random CD structure that underwent a further structural change to yield a typical amyloid β-sheet CD structure with fibrillar morphology when subjected to a different agitation mode. The kinetic barrier was overcome upon initiating the aggregation in the presence of either the seeds of β-sheet fibrils or sorbitol, an osmolyte. We present a case of generating two polymorphs of α-Syn separated by a kinetic barrier. We argue that kinetic control in amyloid polymorphism operates not only during the de novo formation of different polymorphs, but also in their interconversion. One of the therapeutic strategies is to drive toxic oligomers into the formation of relatively non-toxic fibrils. In the same line, sequestration of metastable kinetically trapped supramolecular structures into more stable, non-toxic fibrils could potentially be an important addition to the future roadmap in combating synucleinopathies.

## 4. Materials and methods

### 4.1. Reagents

All plasticwares such as microcentrifuge tubes, 96-well plates, and polypropylene centrifuge tubes were obtained from Tarsons Product Pvt. Ltd. (Kolkata, India). LB agar and broth were procured from HiMedia Labs (Mumbai, India). Sodium phosphate (mono- and dibasic), ammonium sulphate, streptomycin sulphate, ammonium acetate, ampicillin, IPTG, NaCl, and ethanol were purchased from SRL (Mumbai, India). Proteinase K, ethylene glycol, erythritol, sorbitol, and thioflavin-T were obtained from Sigma-Aldrich (St. Louis, MO, USA), and SDS gel system was from Bio-Rad (Hercules, USA).

### 4.2. Protein Expression

The cDNA encoding WT α-Syn was cloned in pT7-7 vector, and was a kind gift from Peter T. Lansbury [83]. The pT7-7-α-Syn construct was transformed in BL21(DE3) *E. coli* strain, plated on LB agar medium containing 100 μg ml^−1^ ampicillin, and colonies were grown overnight at 37°C. A single colony was inoculated in a 5 ml LB medium tube, containing 100 μg ml^−1^ ampicillin, and incubated at 37°C with 200 RPM orbital shaking. The culture was induced with 1 mM IPTG at OD_600_ = 0.8, and was further grown for 5 hours post-induction. A 500 μl culture volume was pelleted by centrifugation at 9000 g, and supernatant was discarded. The pellet was resuspended in 30 μl 1X SDS Laemmli buffer and boiled for 5 minutes. The boiled cell lysate was centrifuged at 9000 g for 5 minutes and supernatant was loaded on 15% SDS-PAGE to check for the protein expression. For purification, the colony expressing the protein was inoculated in 5 ml LB medium and grown at 37°C at 200 RPM shaking. Next morning, 1% of the primary culture inoculum was added to a fresh 500 ml LB medium that was grown at 37°C shaking and induction was carried out at OD_600_ = 0.8. The culture was grown for 4 hours post-induction and was harvested by centrifugation at 9000 g for 10 minutes. The harvested culture pellet was resuspended in 20 ml resuspension buffer (50 mM Tris (pH 8.0) containing 10 mM EDTA and 150 mM NaCl) per gram of culture pellet and was stored at −80 °C till further use.

### 4.3. Protein purification

The protein was purified using the method developed by Volles and Lansbury [83]. Frozen cells (−80°C) were kept in boiling water bath for 20 minutes for lysis. All centrifugation steps during purification were carried out at 9000 g for 45 minutes at 4°C. Subsequently, the lysate was centrifuged. The supernatant was collected in a fresh tube, its volume was measured to which streptomycin sulphate (136 µl of 10% solution/ml of supernatant) and glacial acetic acid (228 µl/ml supernatant) was added and mixed. The solution was centrifuged, and the supernatant was collected, to which an equal volume of chilled saturated ammonium sulphate (1:1 v/v) was added. This was done to precipitate the protein, which was followed by centrifugation. The supernatant was discarded and the pellet was resuspended in 100 mM ammonium acetate and equal volume of ethanol to remove ammonium sulphate and centrifuged and supernatant was discarded. The pellet was resuspended in absolute ethanol to remove ammonium acetate. This suspension was centrifuged and supernatant was discarded. This step was repeated thrice. Pellet in the tubes were air dried to evaporate ethanol. The air-dried pellet was solubilized in a 10 mM sodium phosphate buffer and 100 mM NaCl. The solubilized protein was dialyzed in 2 mM sodium phosphate buffer to remove the small molecules and salts. The buffer was exchanged after 4 hours of dialysis and kept further for overnight. Next morning, the dialyzed protein was filtered through 0.22 μ syringe filter and its concentration was measured by a spectrophotometer at 280 nm using a molar extinction coefficient of 5960 M^−1^ cm^−1^. The protein stock was stored at −20°C till further use.

### 4.4. Aggregation setup

To set up the aggregation reaction, the protein was taken out from −20°C, and placed in a 100 kDa cut-off centrifugation filter (Millipore). It was centrifuged at 3000 g for 10 minutes to separate preformed soluble aggregates from monomeric protein. The concentration of α-Syn in the flow-through thus obtained was measured spectrophotometrically. Finally, the monomeric α-Syn was resuspended in 10 mM sodium phosphate buffer (pH 7.4) containing 100 mM NaCl, at a final concentration of 1 mg ml^−1^. The protein was diluted in aggregation buffer (10 mM sodium phosphate, pH 7.4 with 100 mM NaCl, 0.02% sodium azide, and 20 μM ThT) to a final concentration of 1 mg ml^−1^. A 1 ml solution of the mixture was aliquoted in 1.5 ml microcentrifuge tubes and aggregation was initiated by placing the tubes horizontally on a 37°C shaker-incubator platform (manufacturer name) with a constant orbital shaking of 200 RPM. The diameter of orbital shaking was 25 mm. At different time point, sample was placed in a quartz cuvette of 1 cm path length and fluorescence readings were acquired on a Cary Eclipse Spectrofluorimeter (Varian, Palo Alto, CA). The excitation and emission wavelengths were 445 nm and 482 nm, respectively. The slit width was 5 nm.

In a separate agitation mode, 1 mg ml^−1^ α-Syn in the aggregation buffer was added to a 96 well plate and subjected at orbital shaking in a Varioskan LUX multimode microplate reader (Thermo Fisher Scientific, Waltham, MA, USA; ESW version 1.00.38). To minimize sample evaporation due to the ‘edge effect’, peripheral 2 rows and columns from all 4 sides were left blank, and samples were added only in the central part of the plate. The instrument was operated at 37°C at 900 rpm orbital shaking. Excitation and emission wavelengths were 445 and 482 nm, respectively. The data were acquired online in a kinetic loop with readings acquired every 60 minutes using an excitation bandwidth was 12 nm. Progression of kinetics was monitored online through SkanIt Software 6.1 RE. All measurements were read from top of the plate.

For seeding reaction, the end-stage preformed aggregates were mixed well through pipette and requisite volume of aggregates were immediately transferred to a tube containing fresh α-Syn. To carry out α-Syn aggregation in the presence of sorbitol, a stock solution of 1.8 M sorbitol was prepared in the aggregation buffer (10 mM sodium phosphate, pH 7.4 with 100 mM NaCl, 0.02% sodium azide, and 20 μM ThT). This solution was mixed with a 2 mg ml^−1^ α-Syn protein stock that was already in the aggregation buffer such that the final protein and sorbitol concentrations were 1 mg ml^−1^ and 1.5 M, respectively. The volume of the reaction was made up using aggregation buffer.

### 4.5. Thioflavin-T (ThT) assay

ThT stock was prepared in Milli-Q water, filtered using 0.22-micron Millipore syringe filters, and was stored in dark at 4 ℃. The concentration of ThT stock was calculated using molar extinction coefficient of 24,420 M^−1^ cm^−1^ at 412 nm. α-Syn aggregation was initiated at 37℃ and 200 rpm in the incubator shaker. ThT was added to the reaction mixture at a final concentration of 20 μM. Fibril formation kinetics were followed by monitoring a time-dependent ThT fluorescence of the incubated samples on a Cary Eclipse fluorescence spectrophotometer. The samples were taken out from the incubator shaker in a 10 mm path length quartz cuvette (450 μl volume) and its fluorescence was monitored at 482 nm, with excitation wavelength set at 445 nm. The excitation and emission slit width were 2.5 and 5 nm, respectively.

The kinetic curves measured by monitoring ThT fluorescence were fit to the following equation:

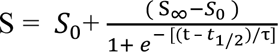

where S_0_ is the spectroscopic signal at time zero, S_∞_ is the final signal, t is the time, t_1/2_ is the time at which the change in signal is 50%, and τ is the time constant. The lag time (t_lag_) was calculated with the equation t_lag =_ t_1/2_ - 2τ.

### 4.6. Atomic Force Microscopy (AFM)

A 10 µl aliquot of protein was deposited on freshly cleaved mica sheets and were allowed to adhere to mica substate for 20 minutes. The samples were extensively washed with 1ml Milli-Q water to remove the salts and osmolytes and allowed to dry for 2 h. Thereafter, AFM measurements were carried out in tapping mode using WITec instrument (GmbH, Germany). Scan speed was 0.5 second/line and 256 lines per images were chosen. The data were analyzed using Project FIVE software package.

### 4.7. Fourier-Transform Infrared Spectroscopy (FTIR)

All FTIR measurements were carried out on a PerkinElmer Frontier FTIR spectrometer (Waltham, MA, USA). The preformed aggregates of the protein samples were centrifuged at 13,000 rpm for 30 minutes. Supernatant was discarded and pellet was resuspended in 200 μl D_2_O, and centrifuged again at the same conditions. This step was performed to remove the residual traces of H_2_O, which interfere in the signal analysis. The centrifuged pellet was removed through a sharp needle and placed on a diamond crystal equipped ATR accessory. The contribution for environmental moisture was corrected prior to sample analysis. For each sample, 50 spectra were acquired and all subsequent data processing were carried out on OriginPro version 9.0. The spectra were averaged and smoothed using a Savitzky-Golay smoothening order 2, and smoothening window of 10. The spectral deconvolution was carried out through a second derivative method and peaks were identified. To identify the area under each peak the peak deconvolution was carried out using peak analyzer tool box of the OriginPro. Briefly, first, the baseline was automatically subtracted from the two end points of the spectrum. This was followed by marking the points on the complete spectra for the peak points that were identified through second derivative method. This step deconvoluted whole spectra into individual peaks with defined peak centres and also revealed the area under each peak. The goodness of fit was ascertained through R^2^ values which was around 0.99 for each fitting.

### 4.8. Circular Dichroism (CD) measurements

CD spectral measurements were carried out on a J-815 spectropolarimeter (Jasco, Tokyo, Japan) equipped with a peltier based temperature controller. The sample was diluted to a final concentration of 0.1 mg ml^−1^ in the respective buffer, and was added to 1 mm pathlength quartz cell. The spectra were collected with data point resolution of 0.1 nm, scanning speed of 100 nm min^−1^ and wavelength range 190 - 250 nm. Three scans were recorded, averaged and plotted. Nitrogen gas flow was maintained at the rate of 5 L min^−1^ and corresponding blanks were subtracted from the respective spectra. The secondary structure deconvolution for each spectrum was carried out through CONTIN and SELCON3 algorithm using DichroWeb software package.

### 4.9. Raman scattering

The native protein and aggregate samples (20 μl) were deposited on a calcium fluoride substrate and were left overnight for air drying. The spectra were collected on a WITec alpha 300 R confocal Raman spectroscopy systems (WITec, GmbH Germany). The instrument was calibrated using a silicon substrate in order to test the standard band position and the intensity. A laser of 532 nm was excited on the sample with a power of 20.2 mW, 600 I/mm grating, and a 100 X plane Fluor objective with numerical aperture of 0.9 at a working distance of 4.7 mm, The data were collected over the range of 100–3,600 cm^−1^, and 10 μ spot site was selected for laser incidence. Cosmic ray and background were removed by filter size 1, dynamic factor 15, and for background removal, shape size 300 and noise factor 100 were taken. Each spectrum consists of 20 accumulations with an integration time of 2 s. After collecting the spectra, all the data were analyzed by origin 9.0 software. Original spectra were smoothened by Savitzky– Golay filter with an order 2 and smoothening window of 10. The smoothed spectra were deconvoluted by 2^nd^ derivative method to locate the sub-bands. The peaks were further deconvoluted by Gaussian Peak Fit module of OriginPro version 9.0. This deconvolution revealed full width at half maximum (FWHM) and area under each peak.

### 4.10. Proteinase K digestion

To 180 μl aggregated sample (1 mg ml^−1^), 0.01 units of Proteinase K was added and mixture were incubated at 25°C. 30 μl aliquot was drawn every 15 min and the reaction was quenched by immediately mixing it with 10 μl 4x SDS-PAGE Laemmlli buffer, and stored at −20°C. Finally, all time point samples were boiled for 5 minutes, centrifuged at 10,000 g, and loaded on 16% tricine-SDS gel. The sample were run at 30 volts for initial 30 minutes and 120 volts for the remaining duration. Thereafter, gel was placed in 10% acetic acid to fix the protein bands. Finally, protein bands were visualized by Coomassie R-250 stain. The band intensity was calculated through densitometry using Fiji image processing package.

## CRediT authorship contribution statement

**Santosh Devi:** Conceptualization, Formal analysis, Investigation, Writing – original draft, Visualization. **Dushyant Kumar Garg:** Conceptualization, Formal analysis, Investigation, Writing – original draft, Visualization. **Rajiv Bhat:** Conceptualization, Writing – review & editing, Supervision, Project administration, Funding acquisition.

## Acknowledgement

Dr. Peter T. Lansbury is gratefully acknowledged for providing α-synuclein DNA clone. Authors thank Jawaharlal Nehru University for infrastructural support. Authors also thank Mr. Saroj Jha, AIRF, JNU for AFM and Raman scattering experiment. Mr. Sandeep Sarpal, AIRF, JNU is acknowledged for FTIR data acquisition. We acknowledge Dr. Rajesh Mishra and Rahamatullah for their initial help in conducting tricine-SDS-PAGE experiments. We acknowledge extramural funding from the Science and Engineering Research Board, Dept. of Science and Technology, Govt of India vide file no. CRG/2021/001396. S.D. thanks Council of Scientific and Industrial Research, Govt. of India for junior- and senior-research fellowship vide file no. 09/263(1213)/2019-EMR-I. D.K.G thanks Indian Council for Medical Research, Govt. of India for research associate fellowship vide grant no. 45/26/2019-BIO/BMS.

## Conflict of interest

Authors declare no conflict of interest.

## Notes

### Competing Interest Statement

The authors have declared no competing interest.

### Summary of Updates

1. Fibril breaking model was included as Fig. 2A and B insets. 2. Fig. panel 2E was replaced. 3. A new Fig. was added as Fig. 7 that indicated seeding of MP aggregation by MP- and SI-derived seeds. The kinetic parameters are included as Table 4.

