## Supplementary material for "Kinetic control in amyloid polymorphism: Different agitation and solution conditions promote distinct amyloid polymorphs of alpha-synuclein": Supporting files.docx

**Supporting Information**

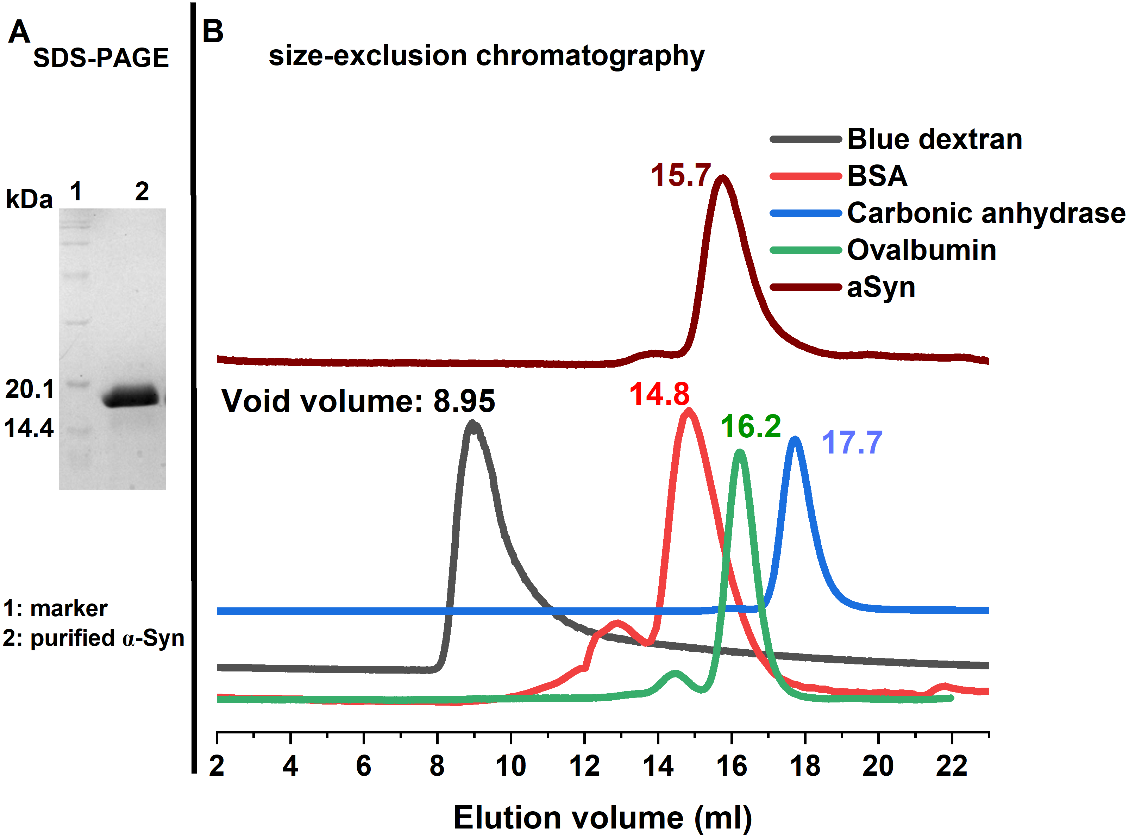

**Fig. S1. Assessing the purity of recombinantly purified α-Syn through SDS-PAGE and size-exclusion chromatography (SEC), as indicated in the figure.** For the SEC analysis, globular protein standards were also run alongside α-Syn.

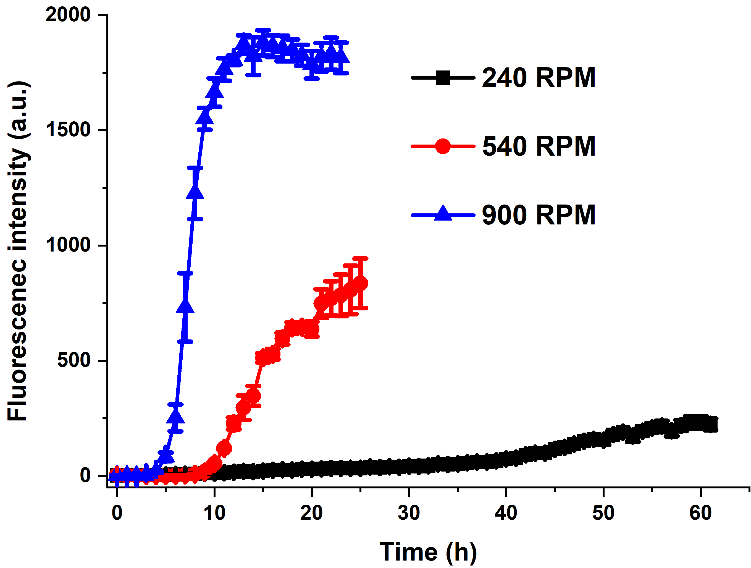

**Fig. S2. Comparing the aggregation kinetics of α-Syn at different agitation speed (as indicated in the figure)**. 1 mg ml^-1^ sample was aliquoted in a 96 well plate and were agitated in a multimode microplate reader.

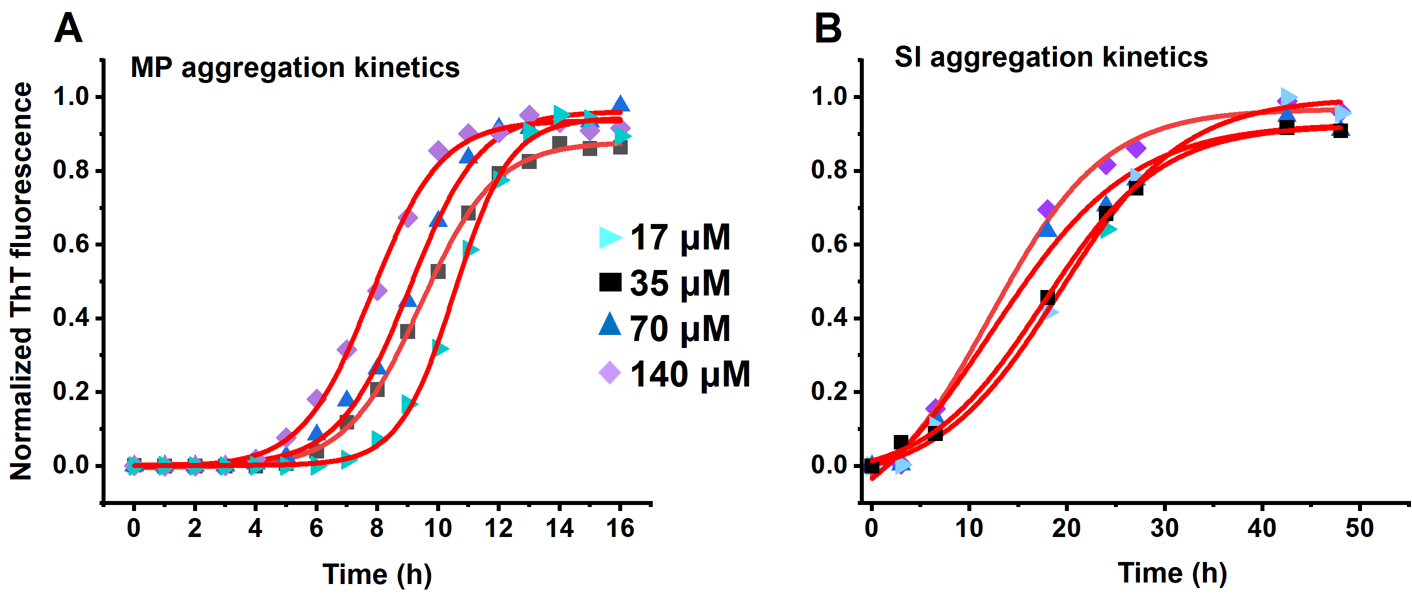

**Fig. S3. Comparing the aggregation kinetics of α-Syn at different monomer concentrations (as indicated in the figure)**.

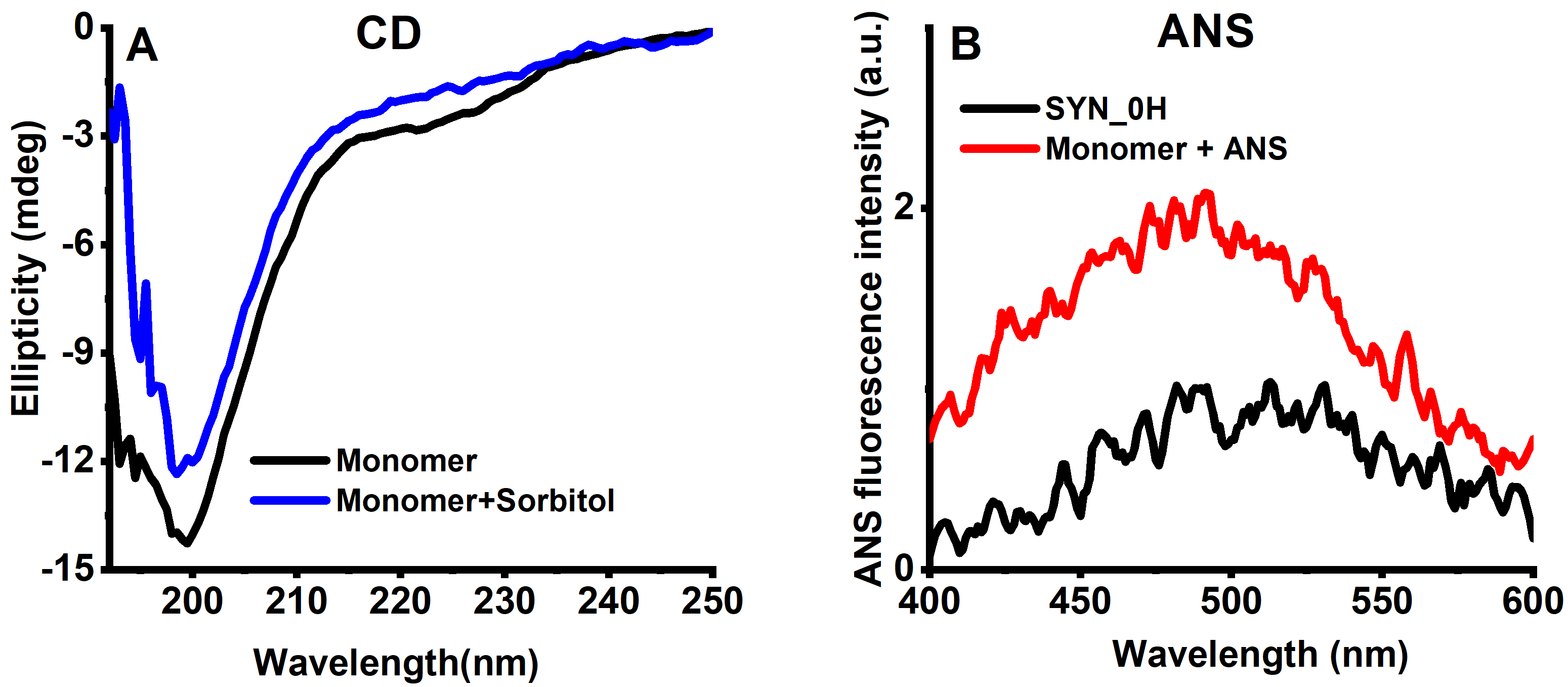

**Fig. S4. Far UV CD and ANS fluorescence spectra of 1 mg ml^-1^ α-Syn with- or without sorbitol.** We observed that the addition of sorbitol did not alter the global structure of α-Syn.

| **Time (h)** | **Software** | **Helix 1** | **Helix 2** | **Strand 1** | **Strand 2** | **Turns** | **Unordered** | **Total** |
| --- | --- | --- | --- | --- | --- | --- | --- | --- |
| **0** | **CONTIN** | 0.008 | 0.028 | 0.1 | 0.054 | 0.081 | 0.728 | 0.999 |
|  | **SELCON3** | 0.022 | 0.02 | 0.03 | 0.027 | 0.036 | 0.822 | 0.957 |
| **5** | **CONTIN** | 0.005 | 0.022 | 0.113 | 0.06 | 0.082 | 0.718 | 1 |
|  | **SELCON3** | 0.035 | 0.045 | 0.082 | 0.048 | 0.084 | 0.682 | 0.976 |
| **10** | **CONTIN** | 0.017 | 0 | 0.05 | 0.025 | 0.014 | 0.893 | 0.999 |
|  | **SELCON3** | 0.034 | 0.03 | 0.037 | 0.043 | 0.032 | 0.83 | 1 |
| **25** | **CONTIN** | 0.009 | 0 | 0.16 | 0.072 | 0.255 | 0.505 | 1 |
|  | **SELCON3** | 0.114 | 0.062 | -0.005 | 0.065 | 0.188 | 0.606 | 1 |
| **50** | **CONTIN** | 0.012 | 0.004 | 0.211 | 0.141 | 0.273 | 0.359 | 1 |
|  | **SELCON3** | 0.012 | 0.013 | 0.317 | 0.161 | 0.252 | 0.252 | 0.989 |
| **80** | **CONTIN** | 0.001 | 0.016 | 0.303 | 0.171 | 0.271 | 0.238 | 1 |
|  | **SELCON3** | 0.001 | 0.023 | 0.285 | 0.176 | 0.276 | 0.229 | 0.99 |

**Table S1A. The relative percentage of different secondary structural elements at different time points during α-Syn aggregation in shaker-incubator (SI) as calculated utilizing CONTIN and SELCON3 secondary structure prediction algorithms in DichroWeb software package.**

| **Time (h)** | **Software** | **Helix 1** | **Helix 2** | **Strand 1** | **Strand 2** | **Turns** | **Unordered** | **Total** |
| --- | --- | --- | --- | --- | --- | --- | --- | --- |
| **0** | **CONTIN** | 0.008 | 0.028 | 0.1 | 0.054 | 0.081 | 0.728 | 0.999 |
|  | **SELCON3** | 0.022 | 0.02 | 0.03 | 0.027 | 0.036 | 0.822 | 0.957 |
| **5** | **CONTIN** | 0.004 | 0.026 | 0.121 | 0.063 | 0.096 | 0.689 | 0.999 |
|  | **SELCON3** | 0.027 | 0.051 | 0.099 | 0.052 | 0.089 | 0.644 | 0.962 |
| **10** | **CONTIN** | 0.001 | 0.025 | 0.167 | 0.09 | 0.117 | 0.6 | 1 |
|  | **SELCON3** | 0.021 | 0.061 | 0.214 | 0.094 | 0.034 | 0.549 | 0.972 |
| **25** | **CONTIN** | 0.003 | 0.019 | 0.149 | 0.071 | 0.088 | 0.67 | 1 |
|  | **SELCON3** | 0.002 | -0.027 | 0.099 | 0.038 | 0.022 | 0.786 | 0.916 |
| **50** | **CONTIN** | 0.003 | 0.016 | 0.162 | 0.082 | 0.122 | 0.615 | 1 |
|  | **SELCON3** | 0.005 | -0.009 | 0.061 | 0.054 | 0.097 | 0.782 | 0.99 |
| **80** | **CONTIN** | 0.004 | 0.022 | 0.16 | 0.09 | 0.122 | 0.602 | 1 |
|  | **SELCON3** | 0.012 | -0.001 | 0.084 | 0.07 | 0.113 | 0.707 | 0.985 |

**Table S1B. The relative percentage of different secondary structural elements at different time points during α-Syn aggregation in microplate shaking mode (MP) as calculated utilizing CONTIN and SELCON3 secondary structure prediction algorithms in DichroWeb software package.**
